# Multi-population dissolution in confined active fluids

**DOI:** 10.1101/2023.09.07.556756

**Authors:** Cayce Fylling, Joshua Tamayo, Arvind Gopinath, Maxime Theillard

## Abstract

Autonomous out-of-equilibrium agents or cells in suspension are ubiquitous in biology and engineering. Turning chemical energy into mechanical stress, they generate activity in their environment, which may trigger spontaneous large-scale dynamics. Often, these systems are composed of multiple populations that may reflect the coexistence of multiple species, differing phenotypes, or chemically varying agents in engineered settings. Here, we present a new method for modeling such multi-population active fluids subject to confinement. We use a continuum multi-scale mean-field approach to represent each phase by its first three orientational moments and couple their evolution with those of the suspending fluid. The resulting coupled system is solved using a parallel adaptive level-set-based solver for high computational efficiency and maximal flexibility in the confinement geometry. Motivated by recent experimental work, we employ our method to study the spatiotemporal dynamics of confined bacterial suspensions and swarms dominated by fluid hydrodynamic effects. Our computational explorations reproduce observed emergent collective patterns, including features of active dissolution in two-population active-passive swarms, with results clearly suggesting that hydrodynamic effects dominate dissolution dynamics. Our work lays the foundation for a systematic characterization of natural and synthetic multi-population systems such as bacterial colonies, bird flocks, fish schools, colloidal swimmers, or programmable active matter.

## 1 Introduction

Active fluids and suspensions comprised of active self-propelling agents such as motile cells or chemically modified colloids are ubiquitous in nature. These out of equilibrium living systems ^1–8^ or synthetic systems ^9–12^ demonstrate fascinating properties such as collective motion, structure formation, intrinsic phase-separation and non-trivial spatiotemporal dynamics such as motility-driven mixing and interface creation ^13–27^. In nature, active systems are sometimes multi-species and often multi-phase, even within the same species. This multiplicity can be the result of distinct species living in the same environment, as is the case in ecological niches, where a heterogeneous mix of cells coexist, and internal boundaries can form, separating cells of two different types. Bacterial colonies, including swarms and suspensions, may feature either symbiotic interactions ^28,29^, or inter-species competition ^30–32^. In bacterial suspensions, chemotaxis and cell death ^19,33^ or the presence of extra-cellular polymers ^20,34,35^ can induce swimming-speed variation leading to phase separation ^19,20^. Equally important are the synergistic effects of motility (self-propulsion at the single cell level), direct cell-cell interactions that include steric and other biophysical or biochemical interactions, and fluid-mechanical inter-cell interactions mediated by the ambient medium. Combined with boundary interactions and confinement, all these can lead to emergent stable spatiotemporal patterns and transitions between states that are structurally distinct; examples of these are swimming-to-swarming transitions demonstrated recently in bacteria confined to wells ^36^, spatiotemporally fluctuating arrayed vortices ^26^, and activity-dependent jamming in growing bacterial colonies ^37–39^. Motility-driven mixing may also result in domain formation, creating interfaces ^21–24^ that may fragment, propagate or mix ^26,27,40^ .

Biological active matter such as bacterial suspensions and swarms are comprised of motile cells that interact with each other, with the ambient medium, and with boundaries. Furthermore, relevant length scales may range from microns (corresponding to single cells), to centimeters or longer (corresponding to system/domain sizes). Experimentally, key parameters influencing emergent dynamics are difficult to control. For instance, controlling cell density, length, and cell self-propulsion speed in bacterial swarms is difficult. Furthermore, investigating how systems change as just one parameter is varied while keeping other parameters fixed is not typically possible. Computations and simulations are, therefore, often used to supplement experiments and probe the effects of parametric variations that are difficult to implement and access via experiments.

Typical computational solutions and numerical simulations follow one of two classes. The first class comprises discrete agent-based methods that directly simulate the dynamics of a collection of interacting agents (cells) in active systems. Such discrete agent-based simulations have been used to study microswimmer suspensions ^41–46^, and active nematic liquid crystalline phases ^47^. Agent-based models without full hydrodynamics (typically called *dry* simulations) were also utilized to study confined single-species swarming bacteria ^36,48^. The second approach focuses on solving mean-field mesoscale continuum kinetic models with coarse-grained interactions. The resulting model may correspond to dry systems without hydrodynamic interactions ^48–50^ or wet systems with hydrodynamic interactions that include a description of the underlying fluid motion ^6,51–60^.

While there has been significant progress in the modeling and understanding of active matter systems, most previous computational studies and models are restricted to single-species or single-phase systems. In many cases, however, biological active fluids constitute two-phase systems or multi-phase systems. This may happen when suspensions or swarms are comprised of a mixture of motile and non-motile cells, cells from the same species but of different phenotypes, or even cells from different species. Indeed, recent experimental work by some of us ^27,40^, and with collaborators ^26^, have elucidated how interfaces between two phases -one active and one passive -evolve in bacterial swarms. Simulating such out-of-equilibrium multi-phase active fluids is challenging. There is clearly a need for general and efficient multi-scale models in multi-species systems.

Motivated and informed by these experiments, we present a mesoscale continuum model governing the evolution of structure and spatiotemporal dynamics in active suspensions and swarms of bacteria. Using adaptive quadtree grids, level set representation, sharp hybrid finite volume/finite difference discretizations, and parallel computations, we formulate a robust, efficient, and scalable numerical approach. Combined with suitable theoretical models, the numerical scheme presented allows for high-throughput simulation and analysis of multi-phase/multi-species systems.

The organization of the article is as follows. In section 2, we first summarize the experimental setup and observations that directly motivate this work on modeling bacterial suspensions and swarms. Governing Fokker-Planck equations are discussed, and steps resulting in the reduction of these high-dimensional equations to moment equations are detailed in the method section. We then present our simulation framework and introduce the parameters that control the spatiotemporal evolution of our computational model system. In section 3, we discuss our simulation results and compare computational predictions with experimental observations. We summarize and conclude in section 4 by providing a roadmap for future explorations using the theoretical framework introduced in this paper.

## 2 Motivations and computational continuum representation

### 2.1 Experiments on bacteria motivate computations

Bacteria growing in colonies, swarms, or suspensions frequently encounter and are confined by boundaries. These boundaries may be hard walls or soft interfaces and lead to strong effects on the collective motion, patterns, and structure formation in these systems. Chen *et al*. ^36^, studied *Enterobacter* swarms confined in microwells and found that self-organized flows comprising a single vortex for small wells and multiple, smaller vortices for larger wells. In previous work, we have demonstrated that swarms of *Serratia marcescens* propagating on wet agar also show similar patterns ^26^, suggesting that these spatiotemporal features are species-independent features. Figure 1(a) shows a representative brightfield image of a *Serratia marcescens* swarm slowly propagating on agar. In the interior of the swarm, intense spatiotemporally fluctuating bacteria and fluid fields develop. Areas within the two red squares were analyzed using cell-tracking combined with Particle Image Velocimetry (PIV) methods; time series of PIV images thus obtained provide evidence of rapidly fluctuating bacterial vorticity and velocity fields (Figure 1(b)). Since the swarm comprises highly motile cells moving in a viscous fluid, we expect hydrodynamic flows generated by collective bacterial motion to also generate fluctuating flow structures with high vorticity gradients.

**Fig. 1.**
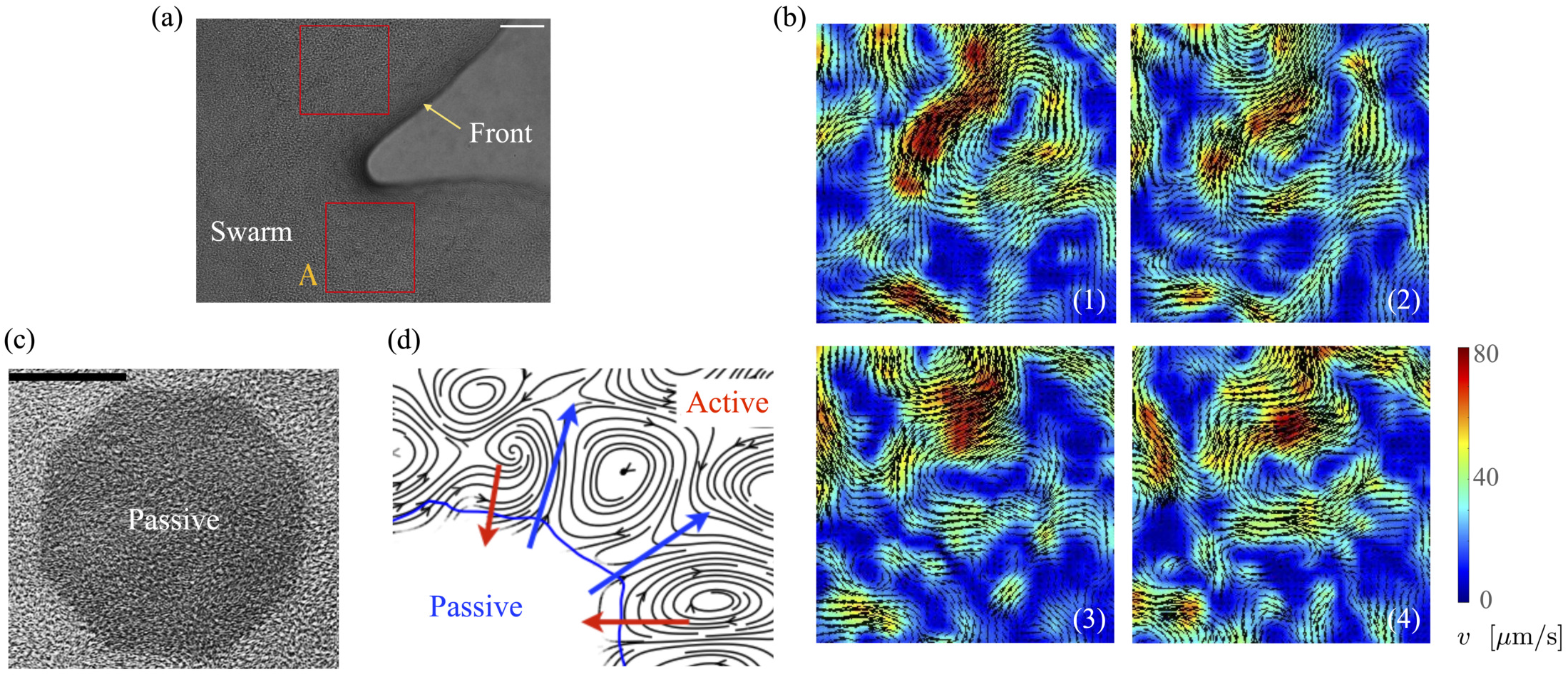
Experimental observations and Particle Image Velocimetry (PIV) analysis of (a) & (b) free swarming *S. marcescens* on agar, and (c) & (d) two-phase active-passive systems. In (a), the colony expands to cover the uncolonized territory. The scale bar is 20 μm. A few bacterial lengths away from the interface, the swarm is well developed with continual and sustained time-periodic flows ^26,27^. We choose boxed regions (50 × 50 μm in area) where these spatiotemporal features are observed for PIV analysis. (b) PIV analysis shows the formation of vortical bacterial flows that are typically 20-30 μm in diameter, with intermittent bursts in the speed of up to 2-3 times this value. (b) Tiles b(1)-b(4) are PIV images in sequence from the red boxed area in subfigure (a); these correspond to time increments of 0.05s. We also plot the magnitude of the bacterial velocity field. Black lines are velocity vectors whose length is proportional to the measured speed. (c,d) Two-phase,active-passive system of a colony generated by immobilizing a sub-domain of the swarm with UV-light (see ^26,27^ for experimental details). The effective area of the passive domain continually reduces as the passive bacteria domain erodes, and bacteria mix with the motile phase.

When a swarm encounters soft boundaries, it may exert stresses and reshape them. To study this, we created passive (immotile) highly frictional domains within active (swarming) domains ^26,27^ by exposing select regions of the swarm to UV light (illustrated in Figure 1(c)). As long as the light is maintained, the passive domain remains compact, and its shape does not change. Upon turning the light source off, the passive domain is continually eroded and fragmented by the invading active phase, eventually mixing completely. The shape of the interface influences the dynamics of the *active two-phase dissolution*. As illustrated in Figure 1(d), corners are flanked on both sides by counter-rotating vortices; the ensuing flow rapidly pinches off the passive phase, yielding further fragmentation and sharpening of the corner. We found that the time for the passive domain to completely dissolve/erode increased with the exposure time τ_exp_ to the UV light, suggesting that the resistance of the passive domain to erosion was dependent on τ_exp_. Furthermore, the temporal evolution of the effective size of the passive region was strongly dependent on exposure times. Large exposure times rendered the passive phase resistant to easy fragmentation.

These prior results strongly suggest that emergent flows driven by activity impact the dissolution process and mixing in multi-phase bacteria systems. What is not clear, however, is the relative importance of these hydrodynamic effects and non-hydrodynamic steric and direct cell-cell effects. Computational modeling of these single-phase and multi-phase systems confers significant advantages in answering this and related questions. First, it is hard to vary activity parameters such as bacterial cell speed and density in experiments. Secondly, while PIV and cell tracking techniques allow us to visualize the collective motion of bacteria, they do not give information on the vorticity and pressure fields in the fluid phase. Both these issues can be addressed in simulations once a suitable model for the system has been identified.

### 2.2 Model and simulation parameters

Readers are referred to the Methods section for derivations. Our mean-field continuum model is an extension of the single population model developed by Saintillan & Shelley ^53,65^. We use the superscript ^(0)^ to refer to the passive population and use the superscript ^(1)^ for the active population. Starting from the Fokker-Planck equation that provides a statistical description of the system in terms of probability distribution functions (Methods, equation 4), the state of each phase (population, with index *i*) is represented by the first three orientational moments of the corresponding probability distribution function. These moments are: (1) the zeroth moment *c*^(*i*)^ corresponding to the local concentration, (2) the first moment **m**^(*i*)^ *i*.*e*., the polarization (polarity) vector that represents the local momentum, with **m**^(*i*)^/*c*^(*i*)^ being the local mean swimming direction (for active swimmers), and (3) the nematic alignment tensor **D**^(*i*)^, that describes the local alignment (Methods, equations 5-7). Scaling appropriately and taking moments, we get dimensionless equations for the three moments (Methods, equations 11-21). Since the two populations are confined to the circular domain and the boundary is impenetrable and non-deforming, we prescribe non-flux boundary conditions on the probability density functions and on the orientational moments (Methods, equations 22-23). The spatiotemporal evolution of the equation for each phase (population) is coupled hydrodynamically to the velocity **u** and pressure *p* fields of the suspending fluid. The fluid is assumed to be Newtonian and incompressible, and therefore, the velocity and pressure fields satisfy the Navier-Stokes equations (Methods, equations 24-25). Finally, these equations are supplemented with the non-slip boundary condition applied to **u** (Methods, equation 26).

Our computational system shown in Fig. 2 consists of a 2-D circular domain Ω of diameter *L* with a rigid and non-deforming boundary. Space in this domain is populated by one of two phases -the active phase (motile cells) and the passive phase (immotile cells). We initialize the active and passive bacteria as two concentric circles, as shown in Fig. 2. For ease of notation, we label the region where the concentration of the passive phase (bacteria) is above the threshold value of 0.5 (see Methods, formal definition in equation (3)) as a crystal. Note that we do not imply that the phase is a crystal in the traditional materials science sense, merely that it is a region where the passive phase dominates. For example, it does not have a repeating crystalline structure, but it tends to align locally and resists the deformation process caused by the active phase.

**Fig. 2.**
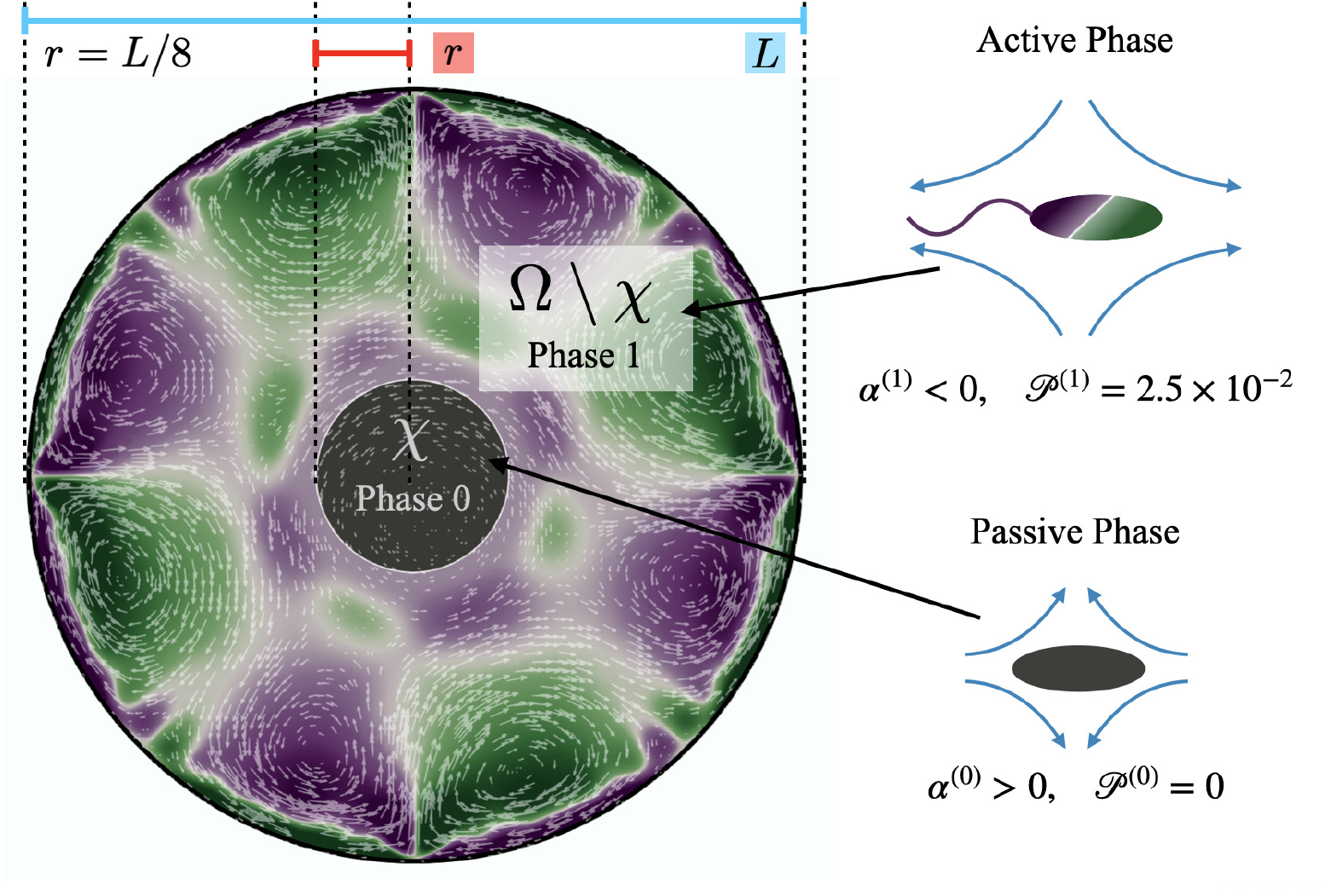
System of interest illustrating the overall setup of the computations and the notations used. The simulation domain Ω is a circle of radius *L*/2. The crystal (passive, grey) domain χ refers to the central circular region that, at time *t* = 0, is comprised of only passive bacteria (contractor, population 0), and the remaining part of the domain surrounding χ is comprised solely of active swimmers (population 1). The fluid is initially at rest, and the populations have no preferred orientation or alignment. As the simulation begins, the populations start aligning, and spatiotemporally patterned vortical flows develop in the active phase. Purple regions indicate positive (counterclockwise) vorticity, and green corresponds to regions with negative (clockwise) vorticity. Color intensity indicates higher magnitudes of vorticity. White arrows indicate the direction and magnitude of velocity. As time progresses, these flows deform, displace, and erode the crystal.

Computations are initialized as follows. Initially, the crystal phase is a circular disk of radius *r* = *L*/8 (Fig. 2, region χ), and the concentration of passive swimmers is set to be 1 in this region. We further assume that all passive cells are contained within this region, and so we set their concentration to be zero outside of χ. The initial active swimmer concentration profile is initialized such that the concentration is zero inside χ and uniform in the annular region Ω \ χ. From this point, we evolve the two-population/two-phase system according to the set of governing equations and numerical approach described in section 4. We track the fluid velocity, concentration, polarization, and nematic alignment of each phase simultaneously. In addition, we follow the evolution of the passive crystal by tracking the boundary corresponding to the threshold concentration of 0.5. This boundary does not have its own special properties such as surface tension or bending energy; it is only an *a posteriori* characterization of the passive concentration.

Our system has four free parameters (see equation 4, and Table 1) for each phase (population). In our simulation, the active and passive phases are comprised of cells with the same geometry. Thus, the two parameters that differ for the two phases are their (swimming) Péclet number 𝓅^(*i*)^, and their activity parameter α^(*i*)^. The activity parameter defined in Table 1 for both passive cells (contractors) and active cells (swimmers) is directly proportional to the strength of the stresslet acting on the fluid around the particle. The passive swimmers are immotile and resist flow deformation. Thus, they have zero Péclet number and a positive *α*^(0)^, which corresponds to a contractile stresslet. Meanwhile, active swimmers are motile pushers and, therefore, have a strictly positive Péclet number. Furthermore, the associated stresslet is extensible, resulting in a negative activity value (*α*^(1)^ < 0). We also assume the cells to have the same translational and rotational diffusivities and thus the same rescaled diffusivity Λ.

**Table 1.** Simulation parameters, non-dimensional groups, symbols, definitions, and typical values. Parameters in bold font represent ones that varied in our simulations.

|  | Parameter | Symbol | Definition | Value | Computational Range | Ref. |
| --- | --- | --- | --- | --- | --- | --- |
| Geometry | Domain diameter | $L$ | | $200 \mu\text{m}$ | | 40,61,62 |
| | Crystal radius | $r$ | | $25 \mu\text{m}$ | | |
| Ambient fluid | Density | $\rho$ | | | | |
| | Viscosity | $\mu$ | | | | |
| | Reynolds number | $\text{Re}$ | $\rho L^2 d_r / \mu$ | $10^{-6}$ | | 56,57 |
| Swimmers | Swimming speed | $V_s$ | | $8.3 - 20 \mu\text{m} \cdot \text{s}^{-1}$ | | 26,63 |
| | Mean density | $n$ | | | | |
| | Rotational diffusivity | $d_r$ | | $0.032 - 2 \text{ rad}^2 \cdot \text{s}^{-1}$ | | 63,64 |
| | Translational diffusivity | $d_t$ | | $0.243 - 100 \mu\text{m}^2 \cdot \text{s}^{-1}$ | | 63,64 |
| | Stresslet strength (Active) | $\sigma_0^{(1)}$ | | | | |
| | Stresslet strength (Passive) | $\sigma_0^{(0)}$ | | | | |
| | Swimming Péclet number | $\mathcal{P}^{(1)}$ | $V_s / d_r L$ | $2.5 \times 10^{-2}$ | | |
| | Passive Péclet number | $\mathcal{P}^{(0)}$ | | 0 | | |
| | Rescaled diffusivity | $\Lambda$ | $d_t / d_r L^2$ | $5 \times 10^{-4}$ | | |
| | **(Active) Swimmer parameter** | $\alpha^{(1)}$ | $\sigma_0^{(1)} n / \mu d_r$ | | $-2\,000 < \alpha^{(1)} < -10$ | |
| | **(Passive) Contractor parameter** | $\alpha^{(0)}$ | $\sigma_0^{(0)} n / \mu d_r$ | | $50 < \alpha^{(0)} < 3\,000$ | |

To reduce the number of free parameters in our computations, we further assume that both populations have the same mean density. Therefore, the variations in the activity levels only reflect variations in the stresslet strength. Because our study focuses on the impact of activity level on the dissolution process, we only vary the activity level and keep all other parameters fixed. For more detailed information on system parameters, see Table 1.

Our model treats individual cells, swimmers, and non-swimmers of any population or phase as rods with defined aspect ratios that can align with the local fluid fields. Steric interactions arising from possible overlap of these rods are not included directly. However, hydrodynamic coupling between the populations via the ambient fluid medium is captured in full. Previous studies have shown that this simpler system reproduces the core dynamics of active suspensions. Our recent experimental work on swarming *Serratia marcescens* suggests that steric interactions and fluid hydrodynamic coupling impact differing aspects of swarming. Specifically, steric interactions impact clustering and aligning. Hydrodynamic interactions also yield alignment but, importantly, intensify and sustain spatiotemporally fluctuating vorticity fields. We expect, therefore, that the dissolution process is dominated by active flow-driven erosion and fragmentation and, therefore, motivates the equations analyzed here.

## 3 Results

### 3.1 Calibration and benchmarks

We calibrate our simulations to match the experimental setup presented in ^40,61,62^, and allow for a meaningful comparison of computational results with experimental data. We start by setting the diameter of the confining geometry to be *L* = 200 *μ*m and the radius of the initial passive crystal domain to *r* = 25 *μ*m. The diffusive properties of the swimmers estimated from the experimental measurements from Travaddod *et al*. ^64^, namely *d*_*r*_ ≈ 0.032 rad^2^/s and *d*_*t*_ = 0.243 *μ*m^2^/*s*. While these parameters are obtained from experiments on *E. Coli*, we expect these values to be representative of other motile bacteria and provide a good value for the lower limit (with higher values corresponding to longer bacterial cells). Bacterial self-propulsion speeds vary across species and also vary according to phenotype. Typically, planktonic cells have slower speeds than swarming cells. From experiments ^26^ on *Serratia marcescens*, we estimate Λ = 1.89 × 10^−4^. Saragosti *et al*. ^63^ estimate *d*_*r*_ ≈ 2 rad^2^/s and *d*_*t*_ = 100 *μ*m^2^/s, and *V*_*s*_ = 20 *μ* m/s, leading to 𝓅 = 5 × 10^−2^, and Λ = 1.25 × 10^−3^. Given the variation in bacteria speeds and thus associated Péclet numbers (see Table 1), we choose to use representative values as follows

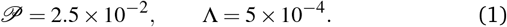

As in our previous studies ^56,57^ we set the Reynolds number to Re = 10^−6^.

The stresslet parameters for the two-populations system, *α*^(0)^ and *α*^(1)^, are hard to estimate *a priori*, primarily because they involve the swimming and contracting stresslet strength, which are hard to access ^40^. Focusing first on active swimmers, we define a meaningful activity range by ensuring that the predominant simulated characteristic flow feature(s) match the ones observed experimentally. As figure 3 illustrates, bacteria are active pushers, and suspensions/swarms of bacteria are destabilized beyond a critical activity level *α*_*c*_. This destabilization leads to the emergence of vortices, whose characteristic size decreases as the magnitude of the activity parameter increases. Following Theillard *et al*. ^56^, the critical activity level can be estimated as

**Fig. 3.**
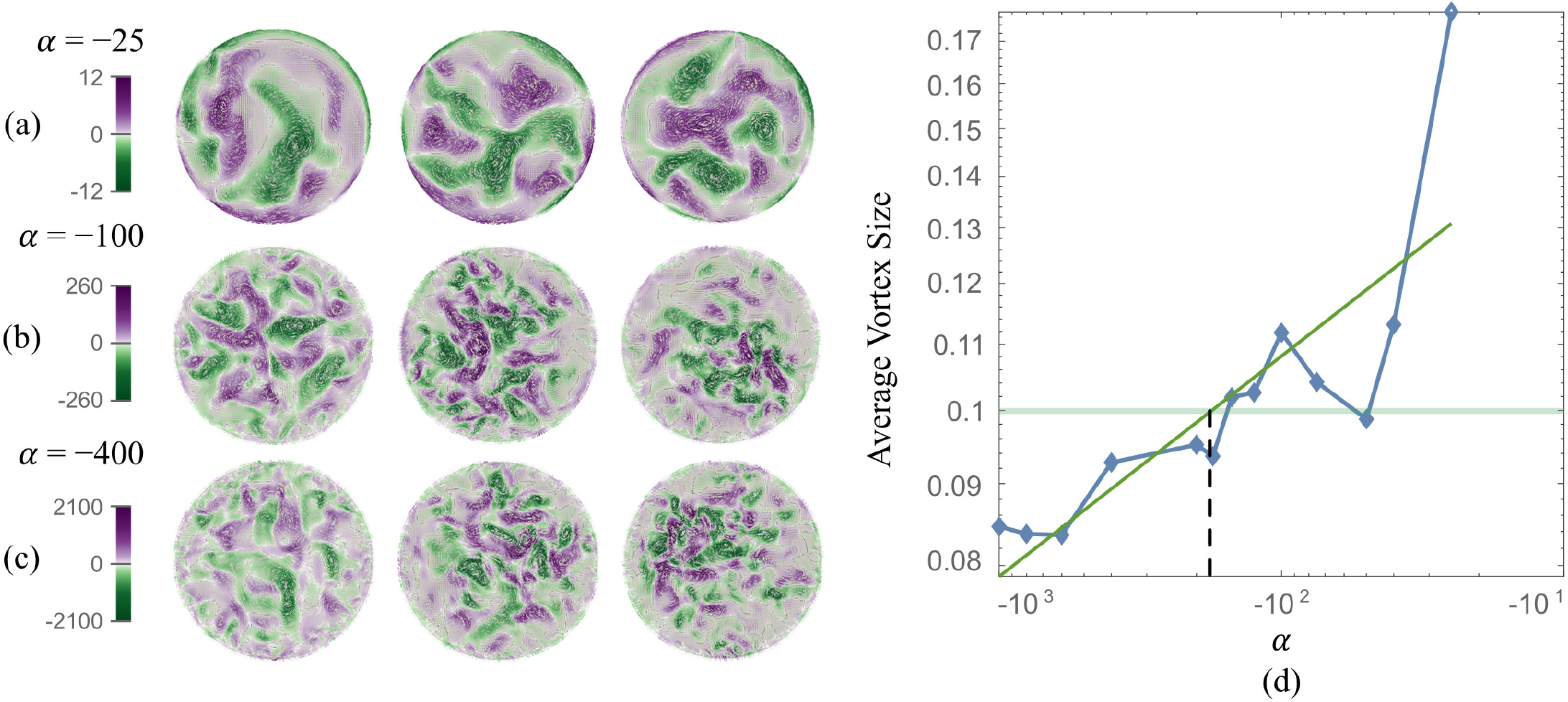
Actively generated vortical flow patterns in single population (phase) suspension. (Left) As the magnitude of the activity parameter |α| increases (from 25 to 400), the generated fluid vortices shrink in size while vorticity intensity (magnitude) increases. For the tiles shown for each series (a)-(c), time is increasing from left to right. (Right) Scaled characteristic vortex length as a function of the activity level (blue diamonds, the solid line serves as a guide to the eye) and power fit (solid green line). The experimental estimation of the vortex size is the horizontal light green line. The vertical dashed line estimates the corresponding activity for our computational model.

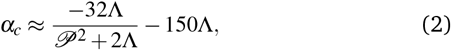

which for the current parameters suggest that *α*^(1)^ must be smaller than −10.07 for the active single-phase suspension to spontaneously destabilize. Indeed, for *α*^(1)^ = −25, −100, −400, we see in Figure 3 the spontaneous emergence of collective vortical flows, indicating destabilization.

In addition, as predicted by the theory, we see the characteristic vortex size shrinking as the swimmers’ activity increases. Swarms are different from suspensions in that the bacteria move near a soft surface, and thus, associated drag frictional forces may be higher. Cell-cell interactions are relatively more important in swarms than in suspensions due to the higher density of cells and longer cell shapes, and hyper-flagellated states attained by swarmer cells. Nonetheless, the same physical argument resulting in instability to activity holds for swarms since the bacteria cells, either singly or in the form of flocks and clusters, effectively act as pushers. Experiments on *Serratia marcescens* ^26,27^, suggest characteristic vortex sizes ∼ 20 *μ*m, which our computations predict is associated with an activity level of *α*^(1)^ ≈ −200. Using this as a point of departure, in our parametric studies, we cover the range −2000 ≤ *α*^(1)^ ≤ −10 to ensure that meaningful characteristic lengths are well-captured. We vary the activity of the passive phase (contractors, *i*.*e*., exerting contractile stresslets) over the range of similar strength 50 ≤ *α*^(0)^ ≤ 3000.

### 3.2 Dissolution dynamics and phase diagram

Having identified physically appropriate ranges of values to use for *α*^(0)^ and *α*^(1)^, we next discuss salient features of the dissolution process and fluid velocity fields that evolve. We formally define the crystal domain *χ* as the area where the concentration of passive particles is above the critical threshold *cT* = 0.5

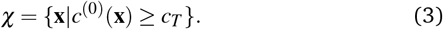

With this definition, we follow the evolution of the shape, size, and properties of the passive domain χ as well as the concentration, velocity, and alignment fields of both populations/phases everywhere in Ω.

Figure 4 illustrates the dissolution process for 100 ≤ *α*^(0)^ ≤ 3000, and for fixed pusher activity *α*^(1)^ = −100 (we recall that the Péclet number is zero for the passive pusher phase). As α^(0)^ increases, we see a stabilization of the crystal domain in that the passive aggregate resists motion, deformation, and fragmentation. Fluid vorticity fields, meanwhile, show interesting features. The maximum fluid vorticity value is only weakly affected, which is expected as it is generated by the active pusher phase with a constant value of α^(1)^. At low α^(0)^, the dissolution process appears to be driven by the mixing of the passive phase by the flow generated by the active phase. For high α^(0)^ values, we see regions of low vorticity appearing around the passive phase. These low vorticity regions are correlated with low concentrations of the active phase. The shape of the crystal domain, along with the associated concentration profile (with relatively low gradients compared to other cases), suggests that diffusion processes drive the dissolution.

**Fig. 4.**
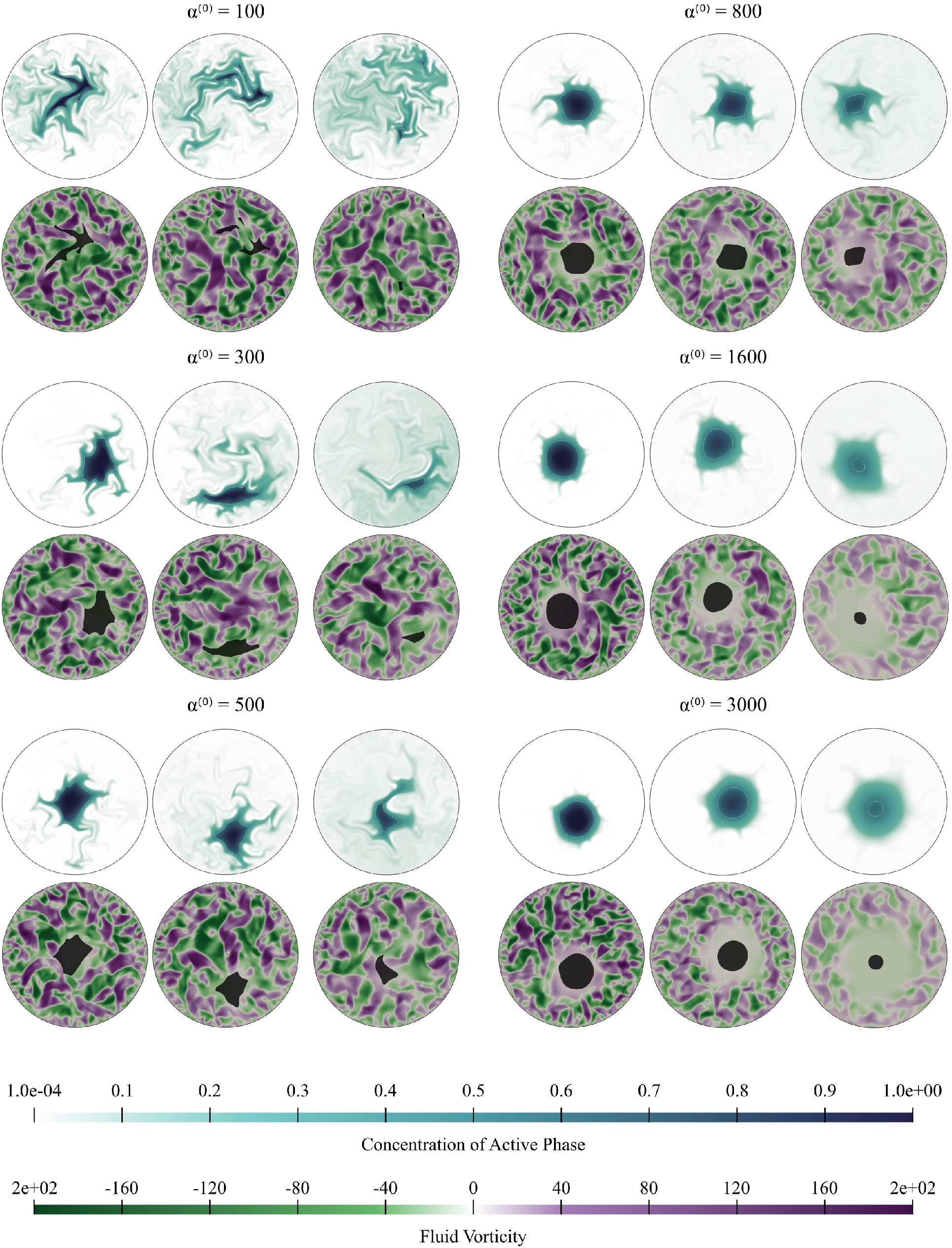
Active dissolution with fixed activity parameter α^(1)^ = −100. We show snapshots of the simulations for increasing value of α^(0)^; effectively, this implies increasing resistance to mixing and dissolution. The passive phase concentration and the fluid vorticity are represented at increasing times from left to right. For α^(0)^ = 1600 and α^(0)^ = 3000, dissolution proceeds slowly with the crystal domain maintaining its nearly circular shape. We also note the exclusion region around the crystal where fluid flow patterns are suppressed. On the other end of the spectrum, for α^(0)^ = 100 and α^(0)^ = 300, mixing drives the dissolution, and the crystal breaks up without suppressing the fluid flow.

The evolution of the passive domain motivates a closer look at the evolution of the crystal domain shape. From a two-dimensional study in α(0) and α_(1)_ parameter space, we compile our results in a phase diagram where we distinguish three different dissolution processes. As we move to the upper right part of the phase plot (for high contractility), the dissolution is predominantly driven by diffusion, and the shape of the crystal domain remains convex throughout the dissolution process. The average curvature of the (crystal) interface is very low, as seen in Figure 7. We define this regime as *convex* (purple circles and region in the phase diagram). For high pusher activity values (high negative values) and α_(0)_ (corresponding to the lower left of the plot), the dissolution process is completely dominated by the flow generated by the active phase. Significant fragmentation is observed, and the passive phase is continually mixed into the fluid. We re-fer to this regime as *fragmented* and depict it in green stars. For parameters corresponding to the transition region between these two regimes, we observe an intermediate crystal dissolution dynamic, which we label as *concave* (blue triangles in Figure 5). This modality is seen when the flow-generating effects of the active phase and dissipating effects of the passive phase are comparable in strength.

**Fig. 5.**
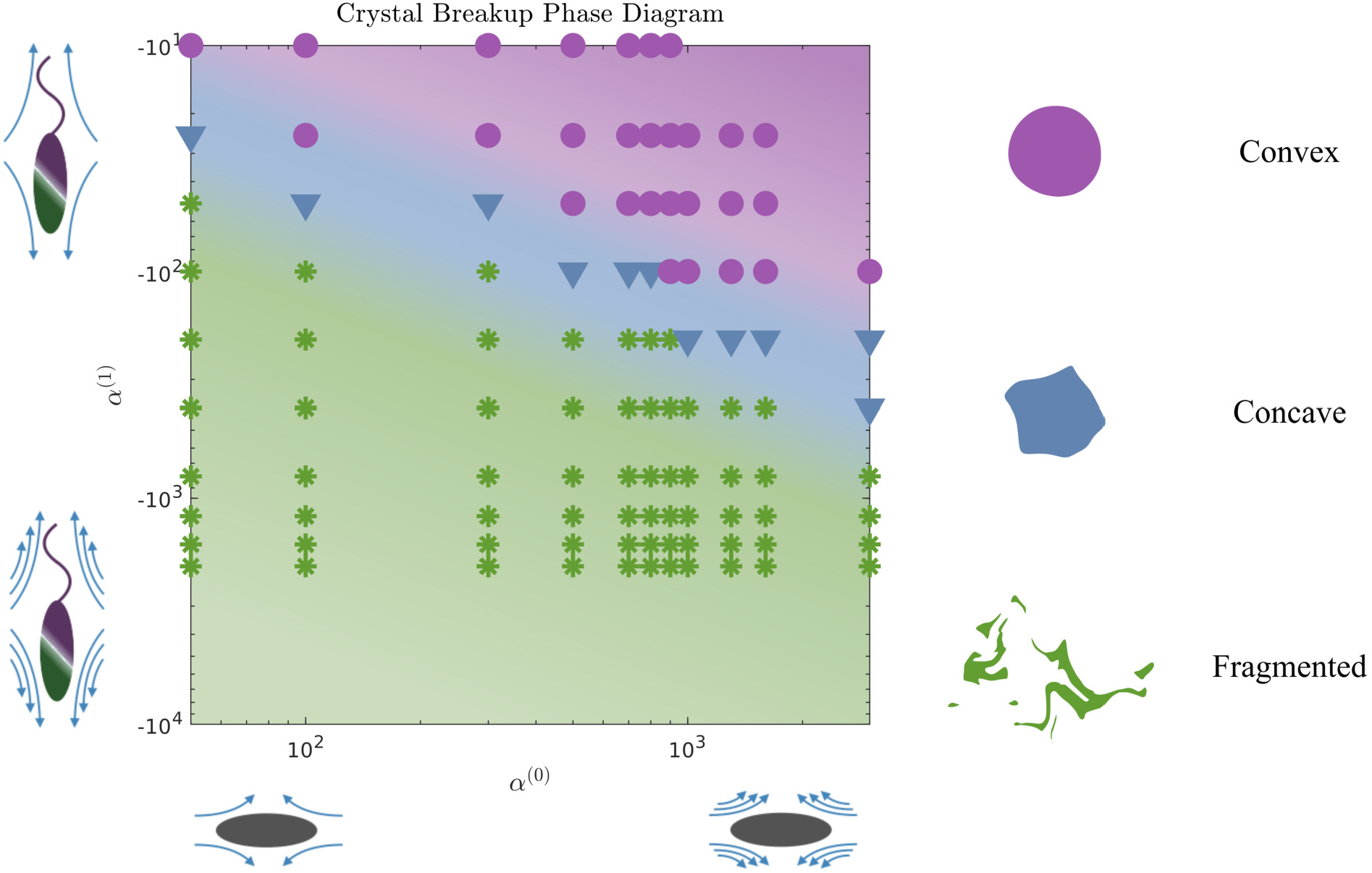
Phase diagram of dissolution modes in α^(0)^ − α^(1)^ space. We identify three different regimes: the *convex* classification (in purple) represents a regime of mostly circular, entirely convex shapes throughout the dissolution process. Comparatively, the *concave* classification (in blue) represents one where the boundary has concave regions as the crystal dissolves. The *fragmented* (in green) classification refers to crystals that break into at least two parts during the simulation, though we usually observe breaking into more than two parts. Active α^(1)^ changes along the vertical axis, increasing pusher activity from top to bottom. Parameter α^(0)^ changes along the horizontal, increasing stresslet strengths going from left to right. Each symbol represents one simulation. The colored background suggests the phase space that maps to the three dissolution modalities.

### 3.3 Quantifying crystal domain shape and properties

We now turn our attention to the evolution of the average fluid velocity, average nematic alignment, the curvature of the crystal, and average vorticity. These variables quantify features of the passive domain, including its shape as well as its connectivity as it dissolves. Thus, we define,

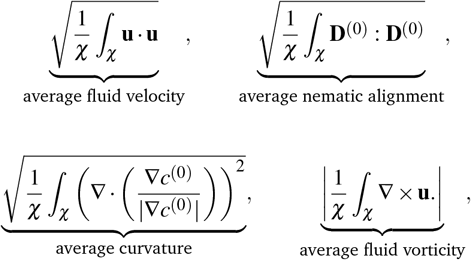

whereas previously stated, χ denotes the area where the concentration of passive swimmer is above the critical threshold *c*_*T*_ = 0.5.

Because the passive cells are immotile, their apparent (lab) velocity is identical to the local fluid velocity **u**. The norm of the nematic tensor **D**^(0)^ quantifies the strength of the local nematic alignment in the passive phase. Averaged over the area χ (that defines the crystal domain), a zero average nematic alignment indicates that the crystal has no preferred orientation. Once the crystal domain is identified, local curvature fields at a point on the interface can be calculated by taking the divergence of the local normal vector. Since concentration fields are used to locate the interface, one can also quantify the roughness of the crystal at any instant in the simulation by evaluating how rapidly the concentration gradients are changing spatially.

Figure 6 displays the averaged curvature of the crystal interface as a function of time for various values of α^(0)^. Time is rescaled by the time at which the passive domain completely erodes and disappears. Thus a scaled time of 0 corresponds to the time when both populations are completely separated, and rescaled time of unity corresponds to when the concentration of the passive phase (bacteria) is less than 50% at any location (*i*.*e*., the crystal has vanished). For α^(0)^ > 800, the average curvature remains fairly constant for the first 80% or so of the simulation before shrinking to zero as the crystal enters its final dissolution stage. This clearly indicates that the crystal remains largely circular and is, therefore, resisting deformation. As α^(0)^ decreases, the passive domain is increasingly stressed and deformed by the active phase and undergoes significant shape changes, which induce large curvature fluctuations. Bursts in curvature are representative of spiky struc-tures on the edge of the crystal (see Figure 6 (a), (b), and (d)). Their short lifespan indicates that these structures are very fragile and quickly disintegrated by the suspension. This behavior is consistent with what is observed in experiments (c.f. Figure 1(d), and ^26^). Interestingly, these fluctuations appear after an initial regime where the curvature remains fairly constant. We understand the presence of this regime as a manifestation of the numerical transient regime, where the diffusive effects are magnified by the initial discontinuity of the concentration profile, and the active fluid flow needs to be established.

**Fig. 6.**
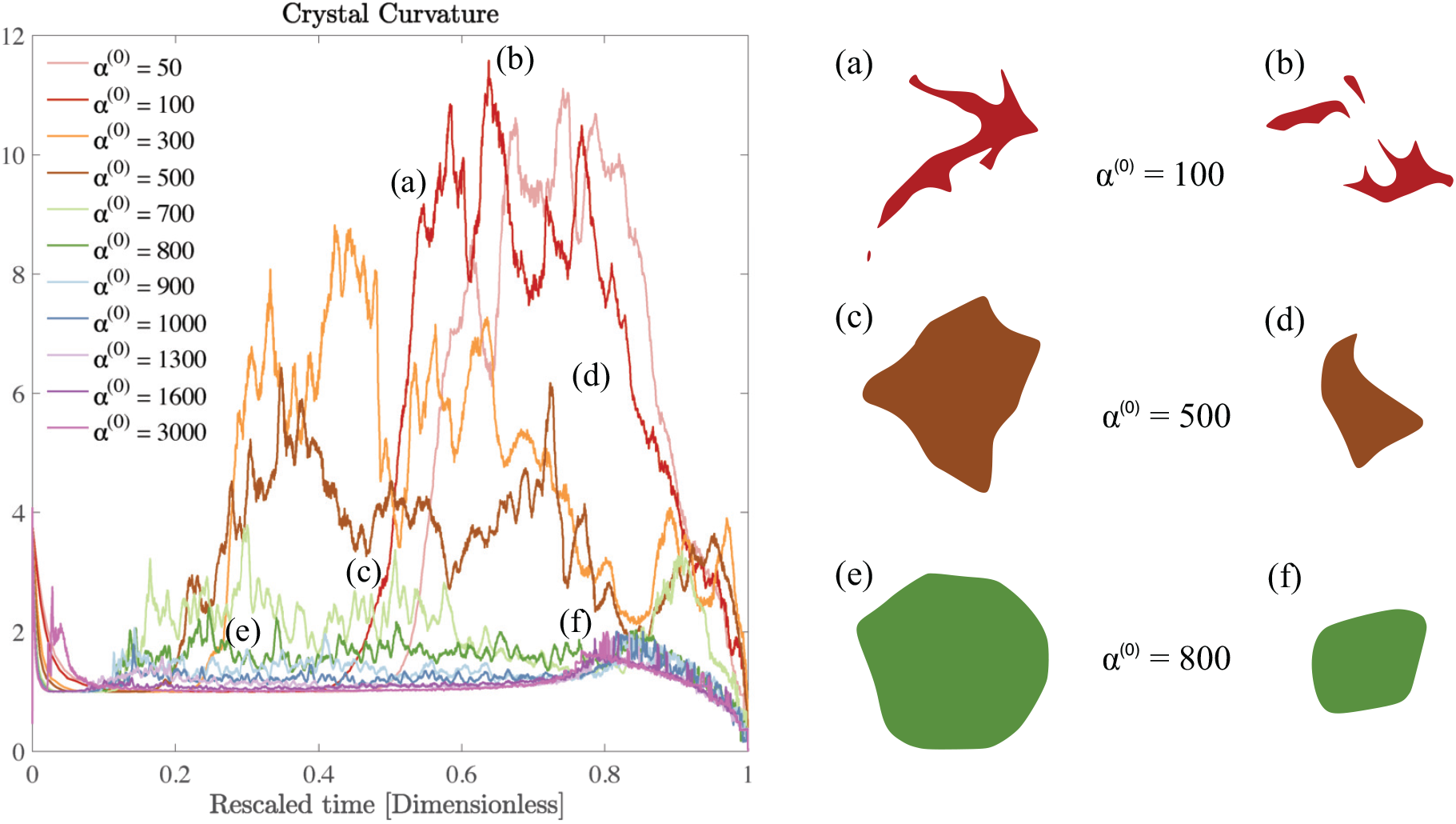
Impact of parameter α^(0)^ on crystal curvature and deformations. (Left)Time evolution of the mean curvature. Note that time is rescaled by the time at which the passive domain completely erodes and disappears. Thus, a scaled time of 0 corresponds to the time when both populations are completely separated, and one corresponds to the first instant where the passive concentration is less than 50% everywhere. (Right) (a)-(f) Instantaneous crystal shapes for selected times and activity levels.

In Figure 7 we display the time-averaged curvature, fluid velocity, nematic alignment, and vorticity for increasing pusher activity parameter α^(1)^. All four quantities are strongly affected by the pusher activity and, as expected, tend to zero as |α^(1)^| → 0. The collapse of the velocity and vorticity curves with α^(0)^ suggests that the crystal average motion and rotation are only driven by the active flows. On the other hand, the curvature and nematic alignment are strongly affected by the contractility for low pusher activity but less affected at high pusher activity. For these two quantities, we interpret the collapse at high pusher activity levels as a consequence of the contractility becoming negligible.

**Fig. 7.**
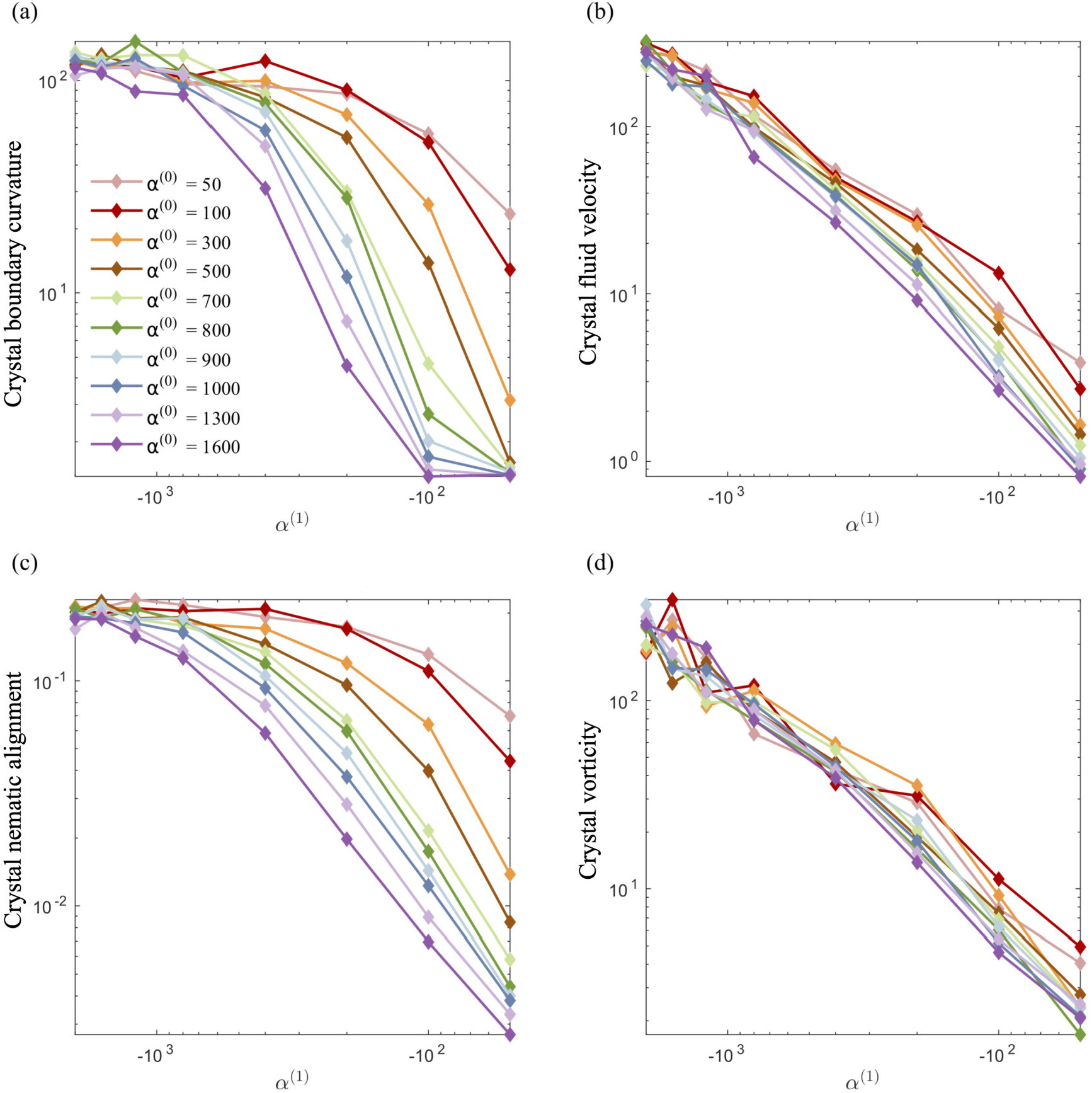
Impact of the activity levels on the crystal average properties: (a) curvature, (b) fluid velocity, (c) nematic alignment, and (d) absolute vorticity. We note that all quantities increase as the strength of the active population increases. The strength has the opposite effect. The observed collapse for (b) and (d) indicates that, for the range of α^(0)^ considered, the strength of the passive population has little to no importance on the crystal displacement or spinning. Conversely, it is essential for the crystal shape and structure, as the spread in (a) and (c) illustrates.

### 3.4 Dissolution times: Comparison with experiments

To map computational results to experimental observations, we first consider the origin of the resistance to deformation in the UV-exposed and unexposed regions of the swarm. The active phase comprises bacteria cells that self-propel through the fluid and exert dipolar stress (extensile stresslets) fields as they move. At the same time, the cells are rigid and cannot deform, and thus, stress fields due to other bacteria are resisted by a contractile stresslet component. Exposing cells to the UV light prevents them from moving *i*.*e*., their swimming Péclet number becomes zero but does not affect their resisting behavior. In the dense swarms, this resistance to induced shear is complemented and accentuated by the collective effects of nearby identically aligned bacteria. In other words, the more aligned adjacent bacteria are, the larger their effective resistance to relative motion and external shear. Since the degree of alignment in the passive region increases with the duration of exposure to UV light τ _exp_, we deduce that experiments conducted at increasing values of τ _exp_ correlate to experiments resulting in higher values of α_(0)_.

Figure 8 is a comparison of experimental data extracted from previously published work ^27^ for dissolution time in swarming *Serratia marcescens*, and a comparison of these with computational predictions. We found that increased exposure time (τ_*exp*_) increases the dissolution time of the passive phase in bacteria-swarming systems. Direct imaging suggested that long exposure times (at intensities beyond a critical intensity) render bacteria in the region immotile ^27^ and result in high values of α^(0)^. In 8(a) and (b), we plot the change in the effective (scaled) radius *r*_eff_(*t*)/*r*_eff_(0) with time for different exposure times τ _exp_. The shrinking of the crystal radius that we compute follows the same qualitative trend as seen in experiments (Figures 8(a) and 8(c)). We understand the overall dissolution process as the superposition of active flow-driven mixing and direct intercalation of active bacteria in the passive phase that corresponds to an effective microscale diffusion process. For α^(0)^ < 1000, mixing is the primary driver of dissolution and erosion. Keeping α^(1)^ fixed (as in the experiments) and increasing α^(0)^ makes the passive domain harder to deform and mix and, thus, to dissolve. Eventually, the crystal becomes too hard to deform, and its dissolution is primarily driven by the diffusive effects. At this point, since the diffusivities are kept constant in our simulations, the dissolution time will start to plateau until the diffusion-only regime is reached. Indeed, in our simulations, we see the dissolution time starting to plateau around α^(0)^ = 1000.

**Fig. 8.**
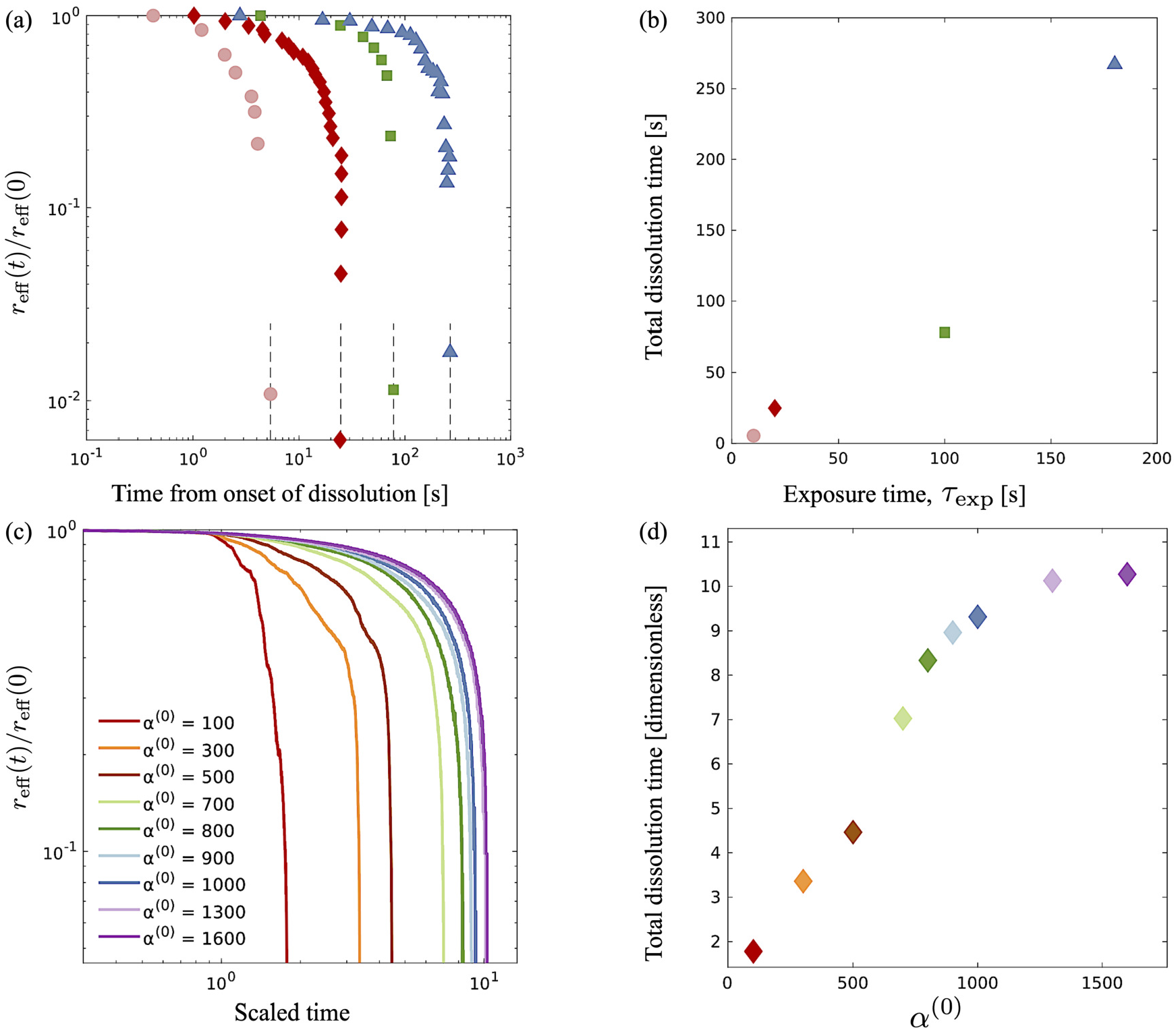
Comparison of experimental data from ^27^ for dissolution time in swarming *Serratia marcescens*, with computational predictions. (a) Evolution of the effective rescaler radius *r*_eff_(*t*) of the compact passive region obtained from ^27^ for various exposure times τ_exp_ = 10, 20, 100 and 180s. The domain size is scaled with *r*_eff_(0), the initial radius. (b) As the exposure time τ_exp_ is increased, the time for the passive domain (crystal, in the terminology used for computations) to erode and dissolve fully increases. These are to be compared with computationally measured effective rescaled radius (c) and total dissolution time (d) for varying levels of contractor activity α^(0)^.

### 3.5 Velocity, polarization, and nematic alignment fields

An advantage of computational modeling for the two-phase system we consider is that parametric sweeps may be used to obtain detailed information for fields related to both the active and passive phases. This is especially true in the case of large α^(0)^ when PIV techniques based on cell tracking fail, and it is difficult to extract local alignment or velocity fields of the passive region. Here, we use our computations to further explore how velocity, polarization, and nematic alignment fields evolve throughout the dissolution process.

Figure 9 displays the net velocity for two values of α^(0)^. At the low value of 300, the net flow in the vicinity of the crystal does not seem to be impacted by the presence of the crys-tal. The net flow surrounding the crystal vicinity strongly fluctuates away from the crystal and also varies greatly over the crystal domain. This results in significant induced interface curvature, strong deformations, and eventual fragmentation and pinching. For α^(0)^ = 800, we see the net velocity being almost uniform in direction and magnitude across the entire crystal, suggesting that the crystal is primarily displaced almost as a solid object. Figure 10 shows the polarization fields corresponding to the snapshots in Figure 9. Again, polarization aligns with the crystal normal at low contractility and, thus, the concentration gradient. The initial active phase concentration gradient near the interface is sufficiently strong to generate fluid flow in the same direction that nematically aligns the active phase, thereby reinforcing the invasion. For low α^(0)^, we also see polarization defect lines running through the crystal (at t = 1.73, for instance). In contrast, for the higher value of α^(0)^, we see the appearance of point “star defects” reminiscent of asters that slowly weaken in magnitude with time.

**Fig. 9.**
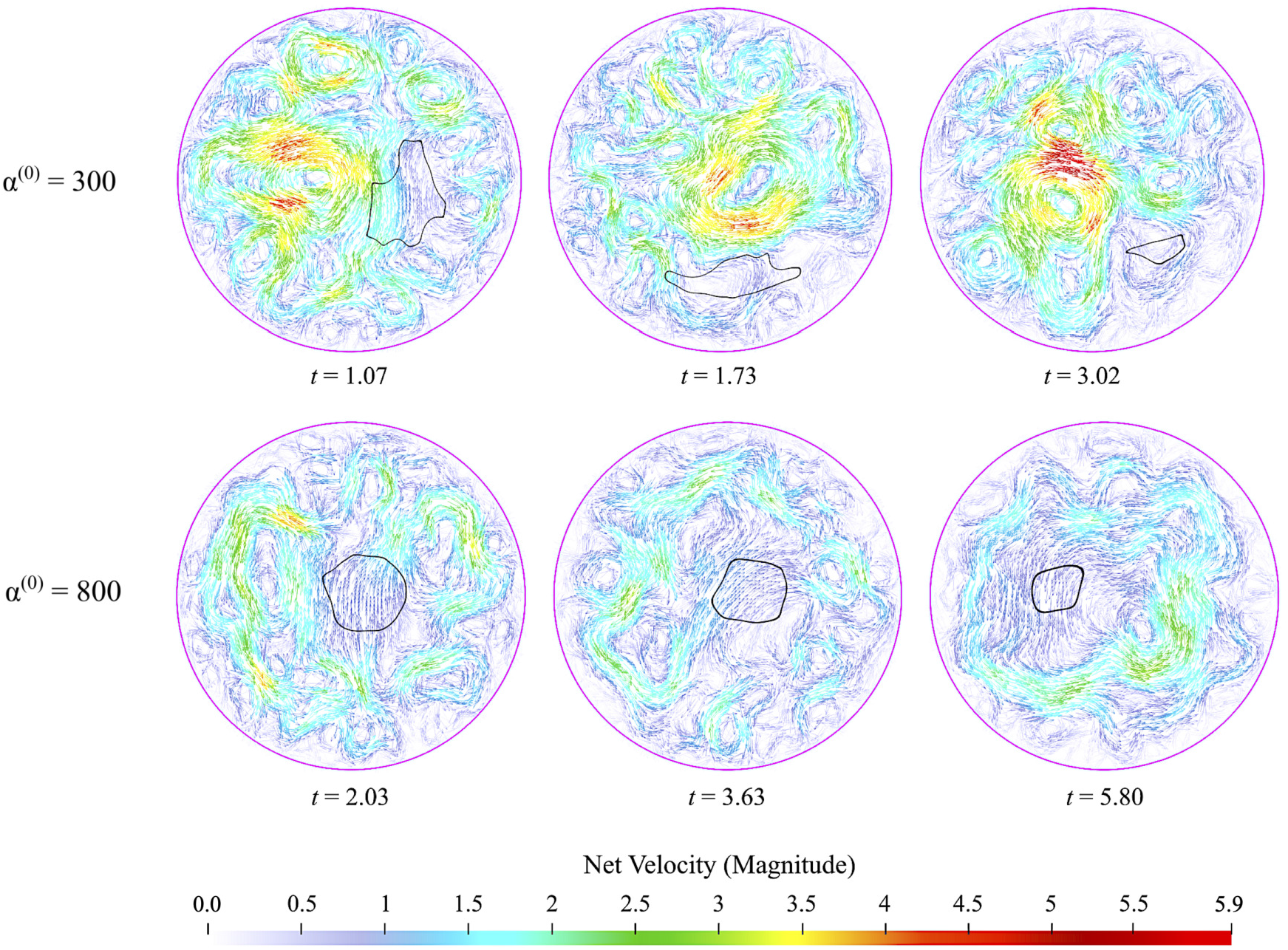
Snapshots of net velocity fields inside the passive domain. The edge of the crystal is indicated as the black curve. Since the passive phase has no intrinsic mobility, the net velocity is the fluid velocity plus the swimming velocity of the active phase. At the lower value of α^(0)^, velocity fields around the crystal domain are strong, and high gradients develop and persist through the entire dissolution process. When α^(0)^ is increased to 800, the net velocity field in the vicinity of the crystal is suppressed significantly, as evidenced by the much lower magnitudes and gentler gradients.

**Fig. 10.**
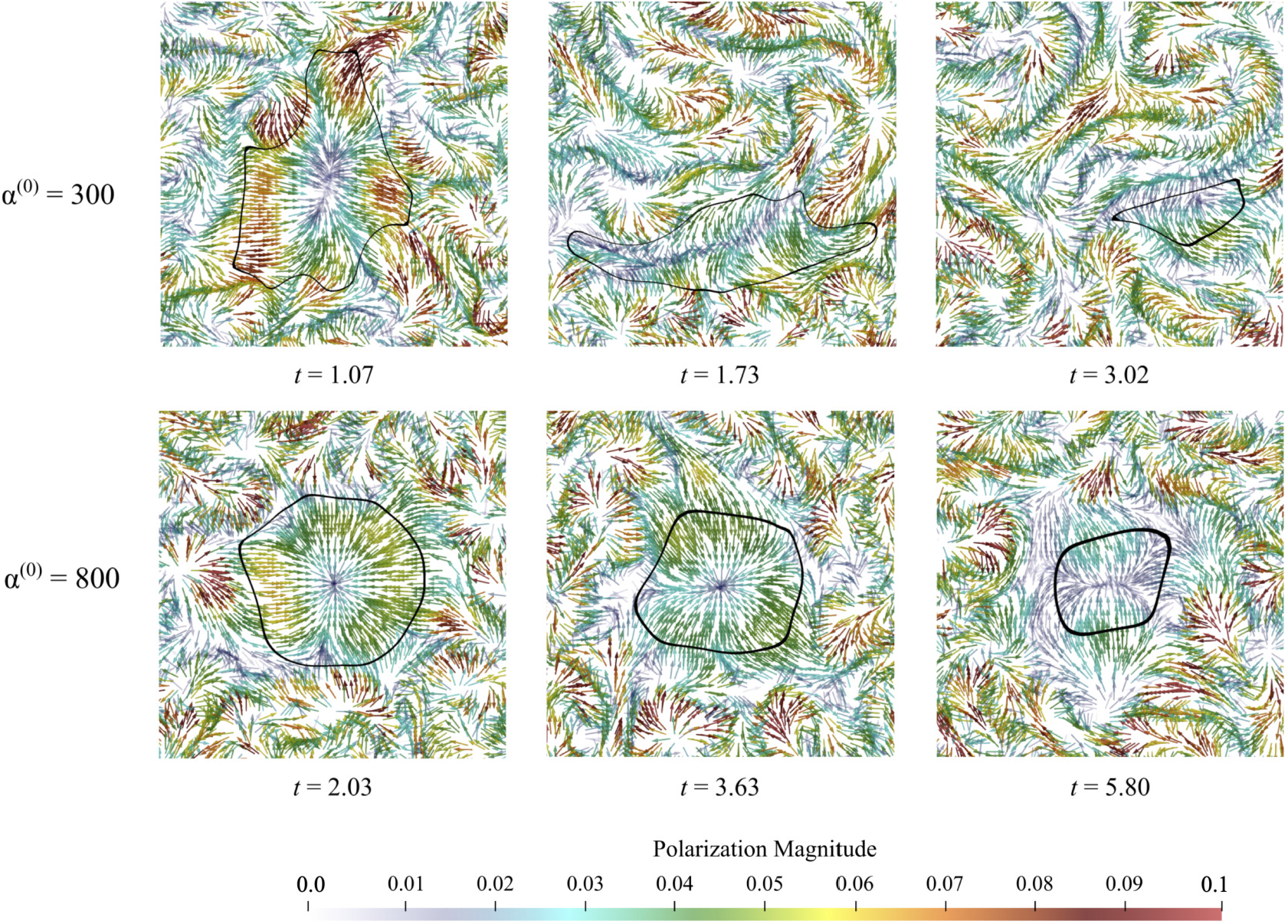
polarization of active phase during the dissolution process. For low values of α^(0)^, This may be because the active phase generates spatiotemporally strong and fluctuating flows that move, deform, and eventually erode and dissolve the crystal with little resistance. Everywhere, lines of strong polarity form and deform continuously, generating highly fluctuating flows and, thus, powerful mixing. For high values of α^(0)^, we see the emergence of star/aster-like defects inside the passive phase, with the magnitude of the local polarization slowly decreasing as the crystal dissolves in time.

In Figure 11, we display the principal direction and corresponding eigenvalue for both populations. Lighter colors show stronger alignment, while darker colors show weaker alignment. These snapshots are consistent with observations made in Figure 4, which showed that the passive phase disrupts flow in the vicinity of the crystal only for high values of α^(0)^. Within the crystal, there are differences between the two experiments. Low contractor (α^(0)^ = 300) activity has higher levels of alignment than high contractor (α^(0)^ = 800) activity. The pusher alignment influences the alignment of the contractors in the α^(0)^ = 300 experiment, and we see regions of agreed alignment inside and near the boundary. For the α^(0)^ = 800 experiment and others in the high-contractility regime, where there is less agreement in alignment inside the crystal and around it. This is because the passive phase slows down the fluid, which drives alignment. When the flow is dis-rupted, the alignment is also disrupted.

**Fig. 11.**
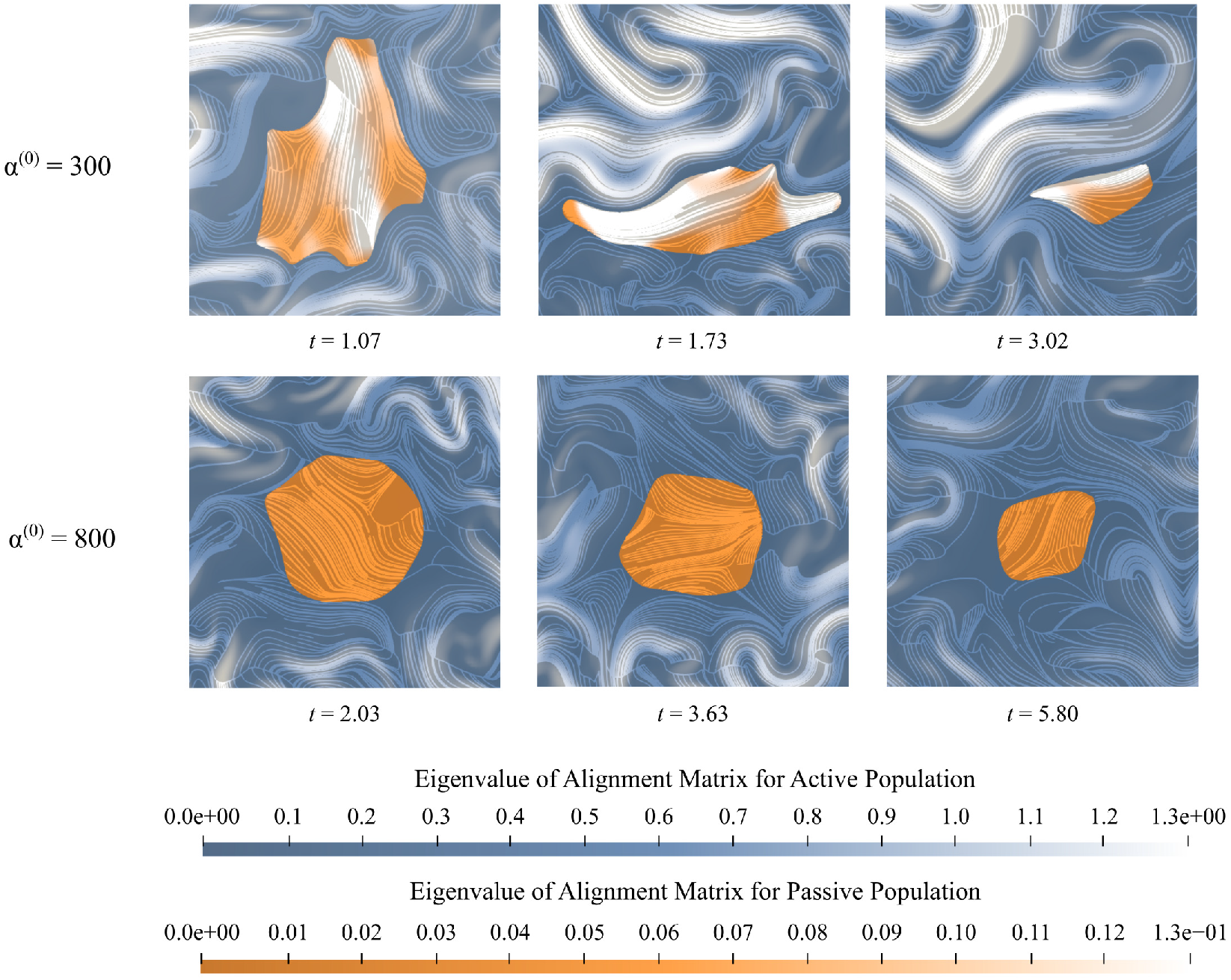
Nematic Alignment -the direction of the streamlines indicates the direction of the eigenvector of D^(*i*)^ associated with the positive eigenvalue. The intensity of colors indicates the relative magnitude of that vector. Lighter colors represent higher levels of local agreement in alignment between the swimmers. Inside the crystal (orange), only the eigenvector and eigenvalue associated with the passive phase are shown, and outside the crystal (blue), only the eigenvector and eigenvalue of the active phase are shown. For low contractor activity, the alignment of the pushers influences the alignment within the crystal, and there are regions of high alignment outside, along, and within the crystal interface. For high contractor activity, there is less agreement in alignment within the crystal, and the alignment strength decreases in the pusher region just outside the interface as the contractors diffuse.

## 4 Summary and discussion

Numerical simulations have played a central role in the investigation of active fluids, allowing one to probe the microscopic mechanisms underlying experimentally observed dynamics. Direct particle simulations of systems such as microswimmer suspensions have been useful for the study of dynamics at small scales. Another approach has been based on continuum kinetic models, which include coarse-grain interparticle and hydrodynamic interactions into systems of mean-field partial differential equations coupling microstructural variables with the underlying fluid motion. All of these models are restricted to single-population suspensions.

In nature, these fluids are usually active multi-phase mixtures. This multiplicity can be the result of distinct species living in the same environment, as is the case in ecological niches, where a heterogeneous mix of cells coexist, and internal boundaries can form, separating cells of two different types. Bacterial swarms and microbes are another good illustrations that can lead to either symbiotic interactions or inter-species competition ^27,66,67^. Recent work conducted on sub-lethal concentrations of antibiotic on swarming and biofilm-forming *Bacillus subtilis* has found that in the presence of sub-lethal concentrations of kanamycin, portions of swarming *Bacillus subtilis* are rendered immotile, forming immotile islands that swarming bacteria must navigate around or dissolve by convecting the immotile cells away ^68^. As a result, the swarm may undergo a phase transition where the immotile islands begin to form increasingly more antibiotic-resistant phenotypes ^39^. Formulating accurate and faithful models that can be applied to explore these systems is challenging, especially as features of interest emerge from the interaction of multiple physics over a wide range of lengths and time scales. There is thus a critical need for efficient, scalable, multi-scale models that can connect the microscale (single-particle level) to the macroscale features and are capable of handling the existence and interactions of multiple species.

Here, we presented a new computational method for modeling such active multi-phase or multi-population systems with hydrodynamic interactions (wet active systems) under geometric confinement. We extended the continuum multi-scale mean-field theory for a single-phase system to multi-phase systems and subsequently analyzed an experimentally motivated two-phase active-passive system. The spatiotemporal evolution of the governing equations for the concentration, polarity, and nematic alignment of each phase (population) is coupled hydrodynamically to the velocity and pressure fields in the suspending fluid. The aggregate material is subject to confining boundaries at which appropriate no-slip and no-flux conditions are applied. The resulting nonlinear set of equations is solved using parallel hybrid level-set-based discretization on adaptive Cartesian grids for high computational efficiency and maximal flexibility in the confinement geometry.

We first validated and benchmarked our code by modeling and reproducing emergent collective patterns observed in confined bacteria suspensions. For this single-population (or single-phase) system, the activity parameter α combined with the degree of confinement determines the size of emergent structures. Keeping the geometry fixed, we find that Lower negative α results in tighter vortexes and smaller characteristic vortex sizes. Based on findings shown in Fig. 3, we identify α^(1)^ = −100 to represent simulated *Serratia marcescens* in the two-phase experiments.

We then analyzed experiments on two-phase *Serratia marcescens* swarms wherein a compact domain of passive immotile bacteria is progressively fragmented and eroded by enveloping active swarming bacteria. These two-phase active-passive systems feature interfaces between the active (denoted by index 1) and passive (index 0) phases that are continuously deformed, fragmented, and convected by local emergent active flows. Our computations capture features of the dissolution process and also predict how dissolution time varies with the resistance of the passive phase to fragmentation and erosion. Our computations confirm that the dissolution process is dominated by the hydrodynamic coupling between the spatiotemporally fluctuating yet strong active flows and the passive domain(s).

Furthermore, we identified two clear modalities by which dissolution occurs. In the first, the crystal (passive domain) remains convex as its area shrinks with time. In the second, the active phase generated flows are strong enough to quickly generate concave shapes and rapid fragmentation into two or more sub-domains, each of which erodes and dissolves. Between these two extremes is an intermediate regime where the crystal holds together in one piece but exhibits concave spots and points of high curvature along its boundary. This is the case that best matches our data from experiments. One of our key findings from our simulations is that compact domains comprised of passive matter can erode/dissolve slowly without fragmentation. The interface between the (outer) active phase and the (inner) passive phase acts as a soft deformable wall that morphs in shape as it reduces in size due to activity-induced dissolution. Our computations predict that by varying the ratio of the activity parameters of each phase, we can select between different dissolution modalities and control the overall time for complete dissolution.

While this study was motivated by fluid-mediated bacterial suspensions and swarms, our approach is generic and is applicable to the analysis of a variety of multi-population systems. Here, we investigated two populations with contrasting activity levels. We are currently studying the impact of internal boundaries and attractive interactions on phase segregation in multi-population systems. With straightforward modifications, our numerical scheme can be adapted to investigate the evolution of nonlinear patterns, such as interacting solitonic waves seen in unbounded active systems ^49^. Further directions would be to study predator-prey systems, excitable matter ^8,25^, or elastically interacting cells ^45,46^. In-depth computational exploration of these systems can yield design principles for engineering applications relevant to soft robotics.

## Methods

### Continuum Modeling of Confined Active Biphasic Suspensions

#### Notations

We consider two active populations suspended in a Newtonian fluid. The whole system is contained in a domain Ω, of characteristic size *L*, and boundary ∂Ω. Each population *i* is composed of Brownian swimmers with mean number density *n*. The swimmers self-propel along their unit vector **p** with constant velocity *V*_*s*_, and have constant translational and rotational diffusivities *d*_*t*_ and *d*_*r*_, respectively. As they swim, they exert a net symmetric force dipole on the suspending fluid with stresslet strength σ^(*i*) 60,69^. We assume the suspending fluid to be incompressible with density ρ and dynamic viscosity μ.

Using *L* as the length scale and 1/*d*_*r*_ as the time scale, for each population, four dimensionless groups emerge from the non-dimensionalization of the governing equations, two groups that involve self-propulsion and activity,

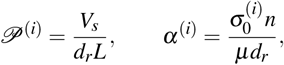

and two quantifying the ratio of diffusive time scales and inertial effects at the scale of the system *L*, respectively,

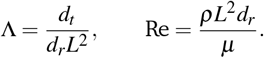

### Moments Expansion

To describe the dynamics of the active suspension, we extend the mean-field continuum model first developed by Saintillan & Shelley ^53,65^ for single-population suspension. In our model, each population is represented by its probability density function Ψ^(*i*)^(**x, p**, *t*), which we approximate by its first three orientational moments *c*^(*i*)^(**x**, *t*), **p**^(*i*)^(**x**, *t*), **D**^(*i*)^(**x**, *t*) using the following closure approximation

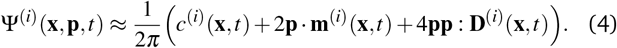

The zeroth moment *c*^(*i*)^ is the local concentration of swimmers. The first moment **m**^(*i*)^, called the polarization, represents the lo-cal swimming momentum and 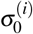 is the local mean swimming direction. **D**^(*i*)^, the second moment in symmetric trace-free form, is called the nematic alignment tensor. It describes the local swim-mer alignment independently of the swimming direction.

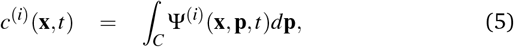

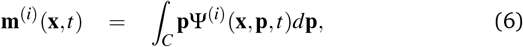

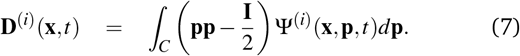

### Governing Equations

The evolution equations for the above moments are obtained by taking the moments of the Smoluchovski equations satisfied by the probability density functions

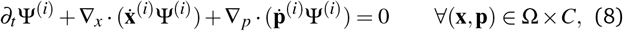

where ∇_*x*_ and ∇_*p*_ are the spatial and orientational gradient operators, respectively. The translational flux velocities 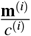 is modeled as

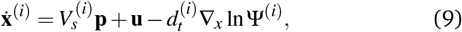

and describes the self-propulsion of the swimmers, advection by the mean-field surrounding fluid flow **u**, and translational diffusion. The orientational flux velocities 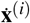 describes the reorientation of the swimmers by the local rate-of-strain and vorticity tensors 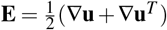 and 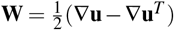 as well as rotational diffusion:

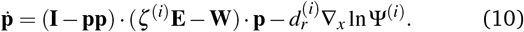

The first term on the right-hand side is known as Jeffery’s equation ^70^, where ζ^(*i*)^, the Bretherton’s constant, is a dimensionless parameter which characterizes particle shape ^71^. ζ^(*i*)^ = 0 corresponds to spheres, such as self-propelled colloids. On the other end of the spectrum, ζ^(*i*)^ = 1 corresponds to slender particles such as bacteria. ζ between 0 and 1 gives corresponding degrees of slenderness. The limits of ζ ≫ 1 and ζ < 0 will be analyzed in a subsequent publication. To obtain evolution equations for each moment, we take the successive of the Smoluchovski equation (8). In dimensionless form, they read

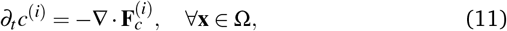

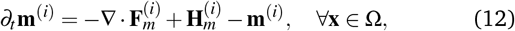

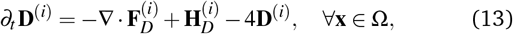

where the translational fluxes are defined as

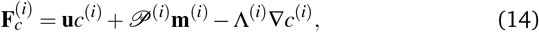

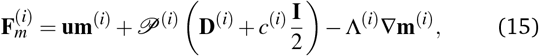

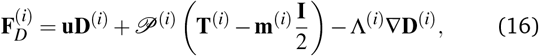

and the hydrodynamic interactions are

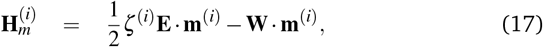

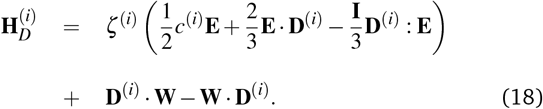

The fluxes 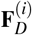 involves the third-order orientational moment **T**^(**i**)^, which we obtain by taking the third moment of the closure approximation (4). Its expression reads

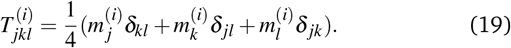

Though not explicitly appearing, the fourth-order moment 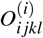 is needed for the equations’ derivation.

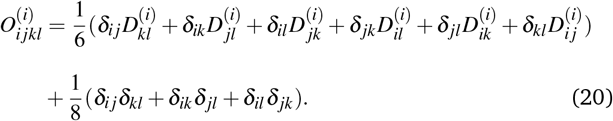

On the confining boundary ∂Ω, we prescribe a no-flux boundary condition on the probability density function ^72^:

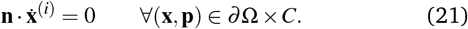

This ensures no swimmers are leaving or entering the domain Ω, and therefore, the total number of swimmers in the system is conserved. It translates into the following no-flux boundary conditions for the orientational moments

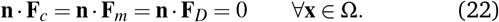

### Fluid Motion

The suspending fluid is assumed to be Newtonian and incompressible, and so the velocity and pressure fields are assumed to satisfy the Navier-Stokes equations

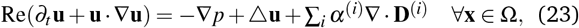

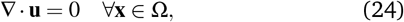

supplemented with no-slip boundary condition

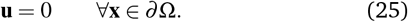

## Numerical approach

The overall numerical method is constructed as an extension of the approach for confined single population active fluids developed by Theillard *et al*. ^57^ and previously used to investigate how the geometry of the confinement can control the emergence of large-scale collective behaviors ^56^. The main difference between the original approach and the one employed for the present study is the ability to handle multiple coexisting populations. The spatial domain is discretized using a non-graded Quadtree grid adapted to the geometry to resolve the thin boundary layers that spontaneously develop. In addition, the grid is dynamically adapted to the spatial variation of the solution: regions exhibiting large gradients in any of the computed quantities. A level-set function characterizes the geometry and can virtually be chosen to be anything.

The numerical solution of the nonlinear system (11)-(12)-(13)-(23)-(24) complemented by the boundary conditions (22)-(25) is constructed using an implicit integrator with hybrid Finite Volume or Finite Difference discretizations. The concentrations are stored at the cell centers for better mass conservation, while all other moments are computed at the vertices for improved accuracy. The fluid velocity and pressure are calculated using the projection-based solver presented by Guittet *et al*. ^73^. We refer the interested reader to our previous work ^57,73,74^ for an in-depth description of the aforementioned numerical techniques and extensive computational validations.

## Acknowledgements

C. Fylling was funded in part by the Center for Cellular and Biomolecular Machines at UC Merced via NSF-HRD-1547848. A. Gopinath and J. Tamayo acknowledge funding from NSF via grant number 2026782 awarded to A. Gopinath. A.T. Standriff made significant aesthetic contributions to Figures 3-11.

## Author contributions statement

**Cayce Fylling**: Formal analysis, Investigation, Software, Validation, Visualization, Writing -Original Draft, Writing -Review & Editing. **Joshua Tamayo**: Experimental investigation, PIV analysis, Visualization, Writing -Original Draft, Writing -Review & Editing. **Arvind Gopinath**: Conceptualization, Supervision, Investigation, Experiments and Analysis, Validation, Writing -Original Draft, Writing -Review & Editing, Funding. **Maxime Theillard**: Conceptualization, Supervision, Formal Analysis, Methodology, Software, Validation, Project administration, Writing -Original Draft, Writing -Review & Editing.

